# Simple spectral transformations capture the contribution of peripheral processing to cortical responses to natural sounds

**DOI:** 10.1101/2019.12.15.877142

**Authors:** Monzilur Rahman, Ben D. B. Willmore, Andrew J. King, Nicol S. Harper

## Abstract

Processing in the sensory periphery involves various mechanisms that enable the detection and discrimination of sensory information. Despite their biological complexity, could these processing steps sub-serve a relatively simple transformation of sensory inputs, which are then transmitted to the CNS? Here we explored both biologically-detailed and very simple models of the auditory periphery to find the appropriate input to a phenomenological model of auditory cortical responses to natural sounds. We examined a range of cochlear models, from those involving detailed biophysical characteristics of the cochlea and auditory nerve to very pared-down spectrogram-like approximations of the information processing in these structures. We tested the capacity of these models to predict the time-course of single-unit neural responses recorded in the ferret primary auditory cortex, when combined with a linear non-linear encoding model. We show that a simple model based on a log-spaced, log-scaled power spectrogram with Hill-function compression performs as well as biophysically-detailed models of the cochlea and the auditory nerve. These findings emphasize the value of using appropriate simple models of the periphery when building encoding models of sensory processing in the brain, and imply that the complex properties of the auditory periphery may together result in a simpler than expected functional transformation of the inputs.

## INTRODUCTION

Sensory systems are biologically very complex, comprising many different structures and cell types that often interact in a non-linear fashion. Furthermore, diverse mechanisms have evolved in different sensory systems for the initial detection, discrimination and encoding of signals at the level of the sensory organs. The complexity of these structures can make understanding the general principles by which they operate challenging. In man-made devices, however, fairly complex circuits may be required to implement straightforward functions due to constraints on their implementation, such as, limited dynamic ranges of the materials, or a requirement of fail-safes. For example, a thermostat is, in essence, a simple negative feedback switch, but in real implementations the circuitry can be substantially more complex because of engineering constraints. Likewise, although neural systems are very complex, under many conditions it could be that the transformations that they compute, at the algorithmic level [1], are substantially simpler than their implementations.

Taking the auditory system as an example, we ask whether the complex mechanisms that characterize processing in the ear have simple algorithmic expressions. Various models of the auditory periphery have been developed over the last thirty years [2–11]. These models have been repeatedly refined in an attempt to account for experimental observations of processing in the cochlea and auditory nerve of a range of animal species and on the basis of human psychophysical data. Some of these models accurately capture the response properties of the auditory nerve and predict their spiking behavior [6,11–16], while others are only a pared down version of the information transformation that might be occurring in the auditory periphery [17–19]. Some of these models have been used to provide inputs for models of cortical neurons [17–20], to generate perceptual models [21], and in machine processing of sounds [2,22]. Few attempts have been made, however, to assess the capacity of different cochlear models in providing input for predicting the response properties of auditory cortical neurons, although some progress has been made in the avian auditory system [23].

Here we consider a wide range of existing models of the auditory periphery, and adapt them to provide input for a phenomenological model of cortical response properties. We also constructed a variety of simple spectrogram-based models, including a novel one accounting for the different types of auditory nerve fiber. We found that primary auditory cortical responses can be explained to a similar degree using simple spectrogram-based cochlea models as input, as when more complex biologically-detailed cochlear models are used. This implies that the complex biophysics of the cochlea may result in a simpler transformation of sensory inputs than this complexity would suggest.

## RESULTS

### Generating cochleagrams using cochlear models

In this study, we consider two broad classes of cochlear models. The first class of cochlear models that we study are based on cochlear filterbanks – these models have somewhat more biological underpinnings. We consider several models of this class, which we refer to here as the Wang Shamma Ru (WSR) model [3–5], the Lyon model [2,10], the Bruce Erfani Zilani (BEZ) [14,15,24] model and the Meddis Sumner Steadman (MSS) model [6,7,11,13,16]. These models vary substantially in their filterbanks and compression functions (see Methods for details). The WSR model has logarithmically-spaced filters, followed by non-linear compression, lateral inhibition and leaky integration [25]. The Lyon model has a log spacing of frequency channels, but this spacing becomes substantially linear near the low frequencies. The frequency decomposition is accompanied by an adaptive gain control mechanism that acts as the compression function [2,10]. The BEZ model includes multiple detailed stages of signal transformation to mimic various stages of the processing by the ear and the auditory nerve of the cat [14,24,26]. The MSS model is similar to BEZ model in that it also models the processing stages from the ear to the auditory nerve, but of a different species, the guinea pig [6,7] (Fig. 1A-C).

**Fig 1:**
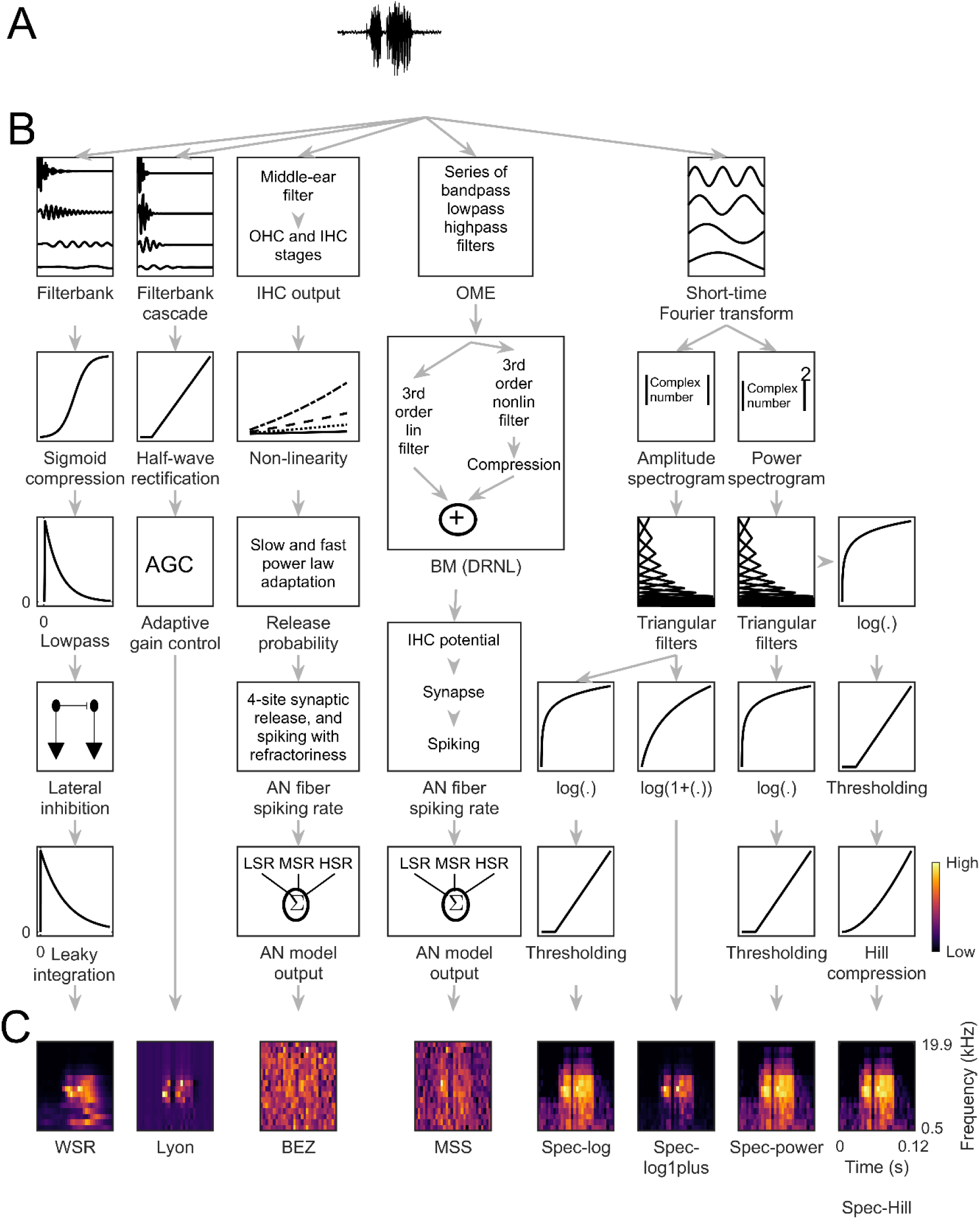
Schematics of the cochlear models. A. A sound waveform, the input to a cochlear model. B. The stages of transformation of sound through each of the cochlear models (from left to right): Wang Shamma Ru (WSR) model [3–5,25], Lyon model [2,10], Bruce Erfani Zilani (BEZ) model [14,15,24], Meddis Sumner Steadman (MSS) model [6,7,11,13,16], spec-log model, spec-log1plus model, spec-power model and spec-Hill model (see Methods). OME, outer and middle ear; OHC, outer hair cell; IHC, inner hair cell; BM, basilar membrane; DRNL, dual resonance non-linear filter; lin, linear; nonlin, nonlinear; AN, auditory nerve; LSR, low spontaneous rate; MSR, medium spontaneous rate; HSR, high spontaneous rate. C. The output of the cochlear models, the cochleagram.

The second class of models are the STFT (short-time Fourier Transform) spectrogram-based models – these models are aimed at approximating the information processing in the auditory periphery without modelling the detailed biological mechanisms. Implementation of these models consists of three key components: frequency decomposition, response integration and compression. We constructed the spectrogram-based cochlear models by performing frequency decomposition using a short-time Fourier transform of the sound waveform. The amplitude or power spectrogram was then put though a weighted summation using overlapping triangular filters spaced on a logarithmic scale to obtain specified numbers of frequency channels. Finally, a non-linear compression function was applied for the amplitude spectrogram-based models. The compression functions used were a thresholded log function, and a log(1+(.)) function. We refer to these models as the spec-log and spec-log1plus, respectively. For the power spectrogram-based models, a thresholded log compression function was used either alone or together with a Hill function; we refer to these models as the spec-power and spec-Hill models (Fig. 1A-C).

Each cochlear model produces a characteristic cochleagram for the same sound input. We illustrate this by presenting a range of synthetic and natural sound inputs to each model. Figure 2 shows the cochlear models’ responses to a click, pure tones of 1 kHz and 10 kHz, white noise and a natural sound (Fig. 2A). For a click input, cochleograms produced by spectrogram-based models have sound energy localized in time, but cochleagrams produced by filterbank-based models are temporally spread, with the response persisting after the impulse occurred (Fig. 2B). For pure-tone input, cochleagrams produced by all models look similar except for the Lyon, BEZ and MSS models, where the cochleagram is broader in frequency content than in the other models (Fig. 2B). For white noise, most models have response distributed across frequency except for the WSR model. Cochleagrams of natural sounds produced by each cochlear model look qualitatively different. However, two filterbank-based models (the BEZ and MSS model) produce similar looking cochleagrams, as do three spectrogram-based models (spec-log, spec-power and spec-Hill). Overall, the WSR model produced very different cochleagrams across a range of stimuli (click, white noise and natural sound). Furthermore, the maximum energy in the cochleagrams of spec-log1plus model is lower than other spectrogram-based models. We quantified the similarity between the cochleagrams produced by each cochlear model for natural sound inputs by calculating the correlation coefficients between the cochleagrams produced by each possible pair of cochlear models (Supplementary Fig. 2.1A). This quantitative analysis supports our qualitative observations (Supplementary Fig. 2.1A-C).

**Fig 2:**
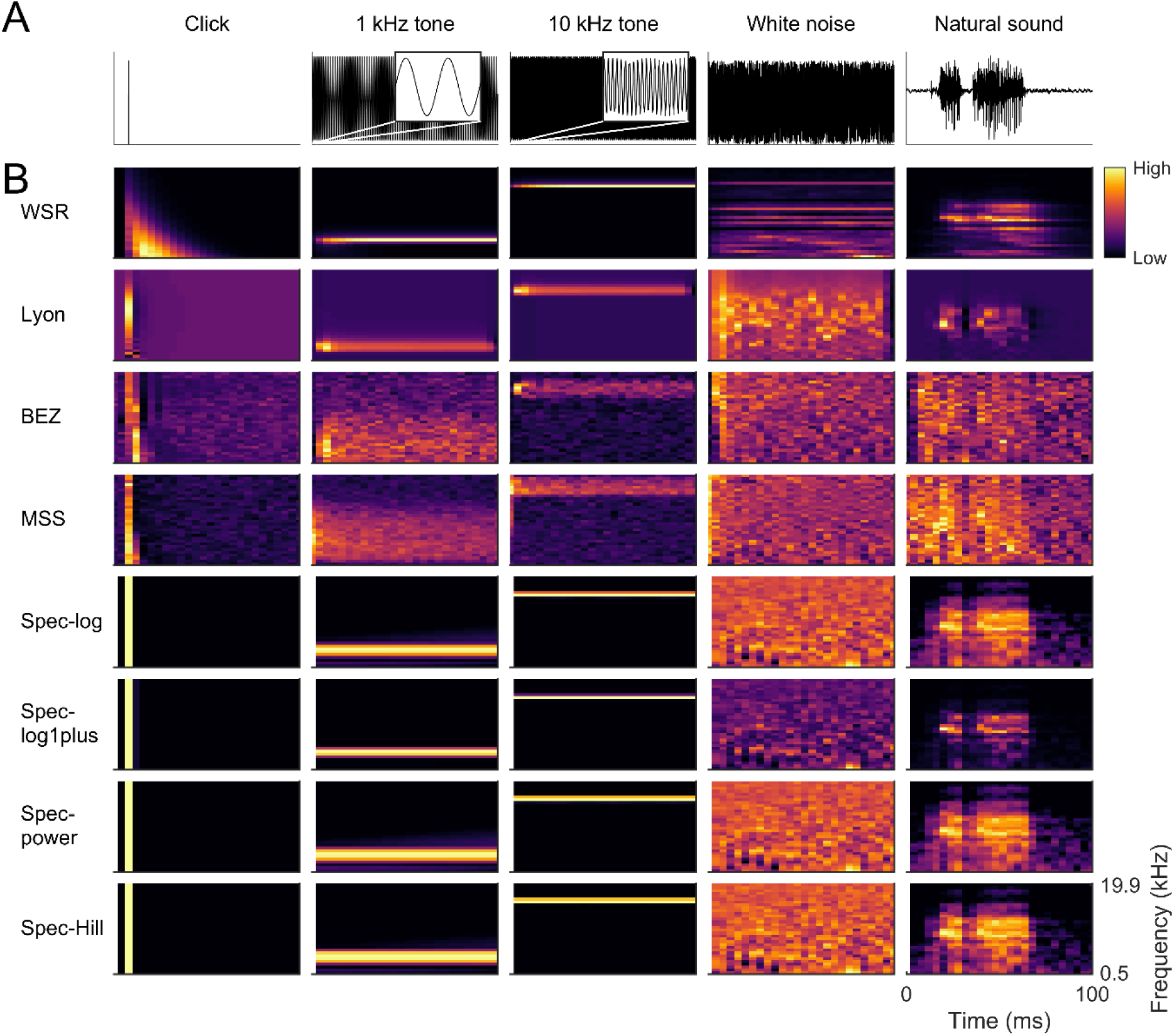
Cochleagram produced by each cochlear model for identical inputs. A. Each column is a different stimulus: a click, 1 kHz pure tone, 10 kHz pure tone, white noise and a natural sound – a short clip of human speech (from left to right). B. Each row is a different cochlear model.

### Predicting responses of auditory cortical neurons using different cochleagram inputs

The data used in this study were derived from extracellular recordings from the primary auditory cortices of anaesthetized ferrets in response to a diverse selection of natural sounds (20 sound snippets, each 5s in duration), including human speech, animal vocalizations and environmental sounds [17–19]. In total, 73 single units were included (see Methods and reference [17] for details on unit selection criteria), selected because they had a noise ratio <40 [27,28] and therefore represented the most reliable responses in our dataset.

The sound pressure waveform is generally not a suitable input to an encoding model of a neuron in the primary auditory cortex. A better choice of input is typically a frequency-decomposed version of the sound [20,23,27,29–40] that resembles the peripheral processing in the cochlea. Cochlear models are often used as input to models of responses of auditory cortical neurons [17–19,23,28,41], such as the commonly used linear-nonlinear (LN) model of neural responses [28,42]. Hence, we use a two-stage encoding framework to estimate firing-rate time-series in response to natural sounds of neurons in ferret auditory cortex. The first stage of the encoding framework processes the sound stimuli using a cochlear model to generate a cochleagram (Fig. 3A). The second stage estimates the firing-rate time-series as a function of the preceding cochleagram using an LN model (see Methods). An LN model was fitted individually to each unit’s responses to 16 of the 20 sound snippets using k-fold (k=8) cross-validation and L1-regularization [43] (distribution of the values of regularization parameter is shown in Supplementary Fig. 3.1). Details of the cross-validation procedure and parameter estimation have been described previously [17] (also see Methods).

**Fig 3:**
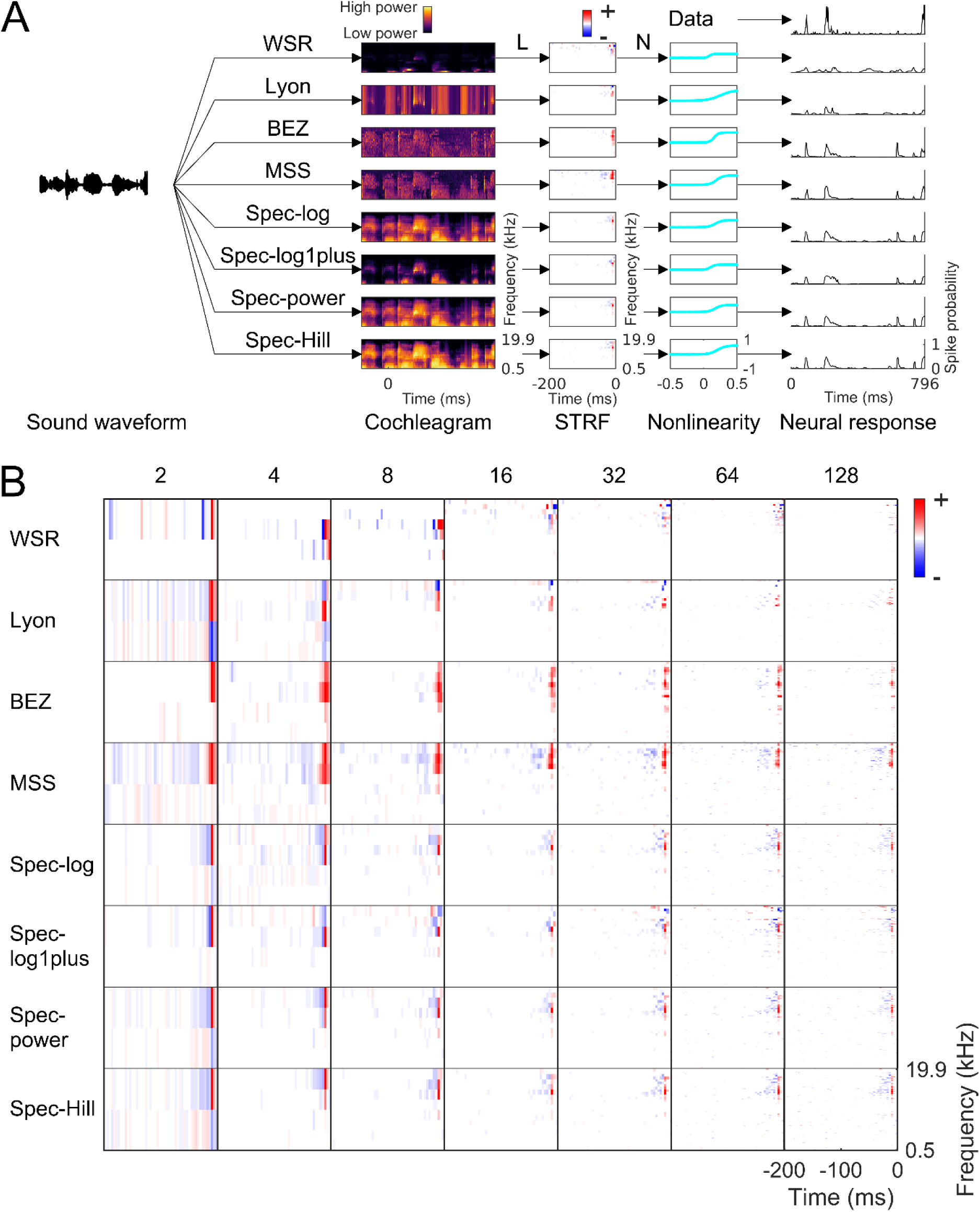
Estimating spectro-temporal receptive fields. A. The encoding scheme: pre-processing by cochlear models to produce a cochleagram (in this case, with 16 frequency channels) followed by the linear (L)-nonlinear (N) model. The parameters of the linear stage (the weight matrix) are commonly referred to as the spectro-temporal receptive field (STRF) of the neuron. Note how the choice of cochlear model influences estimation of the parameters of both the L and N stages of the encoding scheme and, in turn, prediction of neural responses by the model. B. The STRF of an example neuron estimated by using different cochlear models. Each row is for a cochlear model and each column is the number of frequency channels.

The linear part of the LN model captures the linear dependence of a neuron’s firing rate on the frequency content of the cochleagram at different time delays, i.e. the spectro-temporal receptive field (STRF) [20,23,27,29–32,34–40]. STRFs are widely used to describe stimulus feature selectivity of auditory cortical neurons. The general properties of STRFs estimated for same neuron using different cochlear models were similar (Fig. 3B). All cochlear models produced STRFs that contained excitatory and lagging inhibitory fields. The shape of the STRFs produced by different models also resembled each other. The largest weight in the STRF occurred at a comparable frequency (best frequency) and time (latency) for all models and across different numbers of frequency channels. The only exception to this were the 2 and 4 frequency channel models, which sometimes showed very different frequency selectivity, presumably because of the very limited choice of frequency channels (Supplementary Fig. 3.2). The ratio of inhibitory vs excitatory field strength (IE score) was also very similar for STRFs produced by different cochlear models, with the exceptions of the WSR and BEZ models (Supplementary Fig. 3.2).

Although the general properties of the STRFs obtained using different cochlear models were similar, a more detailed analysis revealed some variability between pairs of STRFs estimated for the same neuron using two different cochleagram models (Supplementary Fig. 3.3A,B). Higher correlations were observed between STRFs estimated from same class of cochlear models. In particular, spec-log, spec-power, and spec-Hill models produced very similar STRFs, whereas this was less true of the spec-log1plus model. STRFs obtained with the MSS and BEZ models were similar to each other and, to a lesser extent, to the Spec-Hill model. Applying Gaussian blurring to account for frequency or temporal shifts in the STRFs improves the correlations, but did not change the overall trends in these results (Supplementary Fig. 3.3C).

The prediction performance of the LN model on a held-out dataset also differed between different cochlear models. As a measure of the prediction performance we used the normalized correlation coefficient (CC_norm_) [44] over all neurons in the dataset, where a CC_norm_ of 0 indicates no correlation between the neural response and the model’s estimate and a CC_norm_ of 1 indicates that the model can predict all aspects of the firing rate (averaged over repeats) that depend on the stimulus. We found that the mean CC_norm_ over all neurons varied depending on the choice of cochlear model and the number of frequency channels in the cochleagram (Fig. 4 and Table S4.1). The mean CC_norm_ for the best WSR model (8 channels), Lyon model (64 channels), BEZ model (128 channels), MSS model (128 channels), spec-log model (64 channels), spec-log1plus model (64 channels), spec-power model (64 channels) and spec-Hill model (64 channels) were respectively 0.462, 0.662, 0.644, 0.725, 0.721, 0.630, 0.722 and 0.726 (Fig. 4I and Supplementary Table 4.1). Note that spectrogram-based models, in general, perform better than other models (Fig. 4I). Overall, a log-spaced power spectrogram with successive log and Hill compression functions provided the best prediction performance, with a mean CC_norm_ of 0.726 (Supplementary Table 4.1).

**Fig 4:**
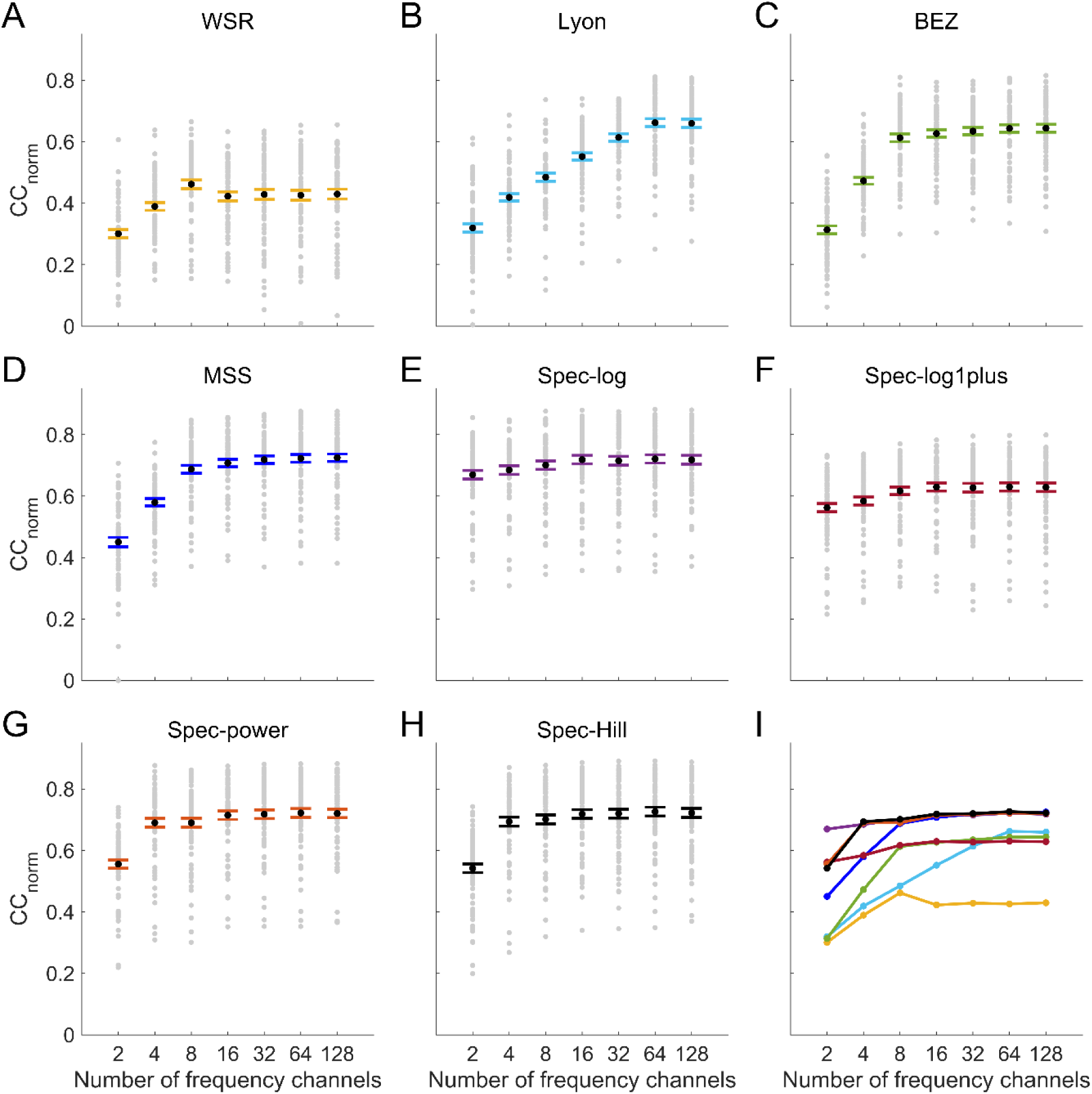
Performance of different cochlear models in predicting neural responses. A. WSR model. B. Lyon model. C. BEZ model. D. MSS model. E. Spec-log model. F. Spec-log1plus model. G. Spec-power model. H. Spec-Hill model. Each gray dot represents the CC_norm_ between a neuron’s recorded response and the prediction by the model; the larger black dot represents the mean value across neurons and the error bars are standard error of the mean. I. Comparison of all models. Each curve is color-coded according to the model it belongs to.

Selecting the best model for individual neurons supports the findings based on the average performance of each model for all neurons. The spec-Hill and MSS models with either 64 or 128 frequency channels provided the best prediction performance for most neurons (Supplementary Fig. 4.2). We also compared the predicted response obtained with the MSS model to the predicted response of the other models, and found similarity in CC_norm_ performance generally co-varied with the similarity in predicted response (Supplementary Fig. 4.3). The Lyon model was an exception to this.

### Multi-fiber cochleagrams

We have used the word cochleagram so far to refer to a time-frequency representation of the sound stimulus that depicts the changes in sound energy across a set of frequency channels over time. In the auditory system, however, afferent nerve fibers tuned to the same frequency can have different thresholds and dynamic ranges [45,46]. To study the impact of this representation on the prediction performance of modeled cortical responses, we used an MSS model with three different fiber types (multi-fiber MSS model) as input to the LN model. Three types of fibers varied in spontaneous activity – either having low spontaneous rate (LSR), medium spontaneous rate (MSR), or high spontaneous rate (HSR). We also constructed a multi-threshold spec-Hill model, where each frequency channel went through three different hill functions with different thresholds and saturation parameters (see Methods). This produced a cochleagram representation that assigns the changing sound level in a single frequency channel into three separate channels analogous to three fiber types in the multi-fiber MSS model.

When we use these models as input to the LN model of cortical neurons, we were able to predict cortical responses to natural sounds slightly better than the single-fiber or single-level versions of the models (Fig 5). The multi-threshold spec-Hill model performed better than the multi-fiber MSS model for cochleagram inputs with fewer than 32 center frequencies, but performs slightly worse for cochleagram inputs with 32 or more center frequencies (Fig. 5).

**Fig 5:**
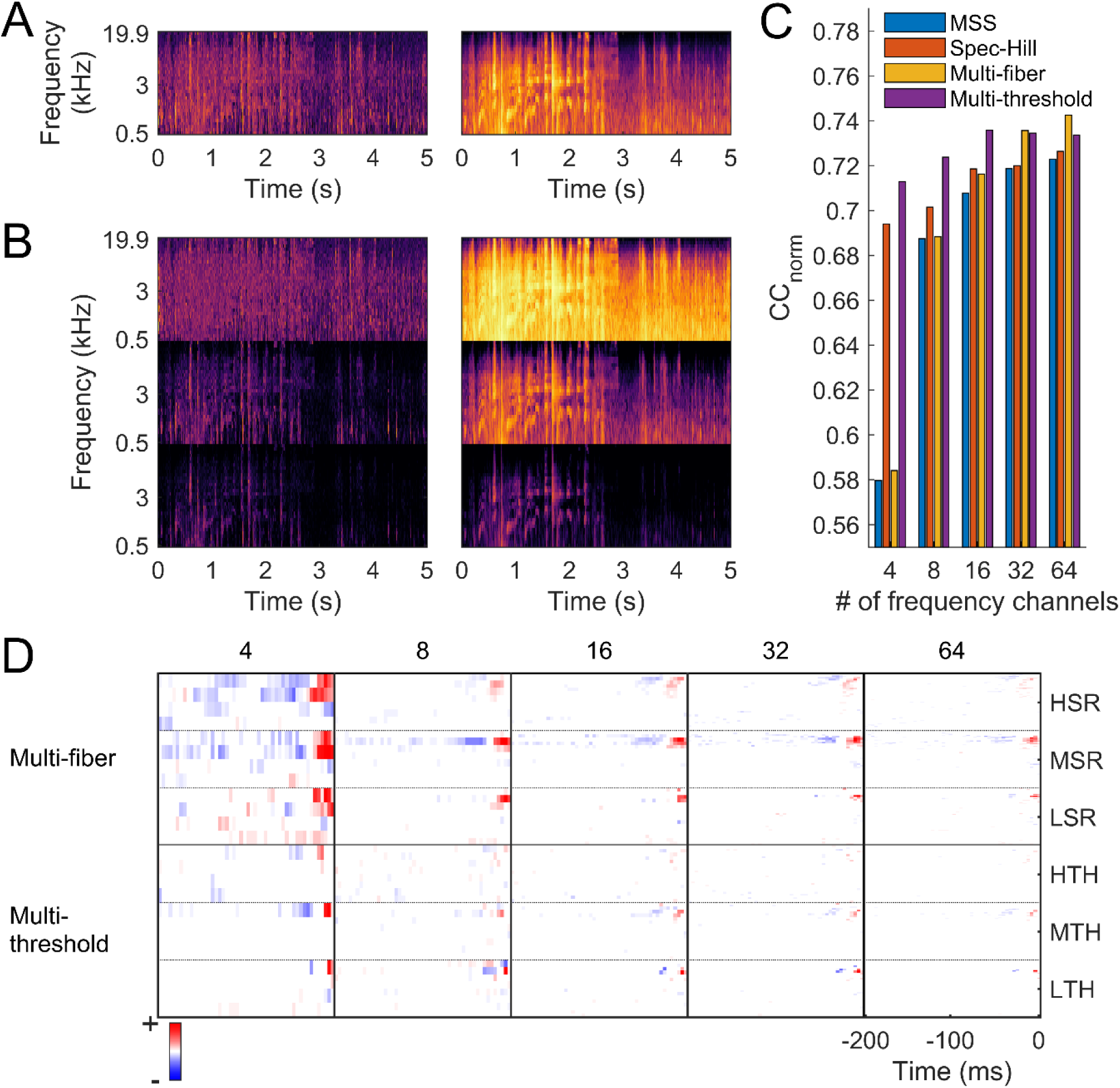
Multi-fiber and multi-threshold cochleagrams. A. Cochleagram of a natural sound clip produced by the MSS model (left) and the Spec-Hill model (right). B. Cochleagram of the same natural sound clip produced by the multi-fiber MSS model (left) and the multi-threshold Spec-Hill model (right). C. Mean CC_norm_ for predicting the responses of all 73 cortical neurons. D. STRFs estimated using the multi-fiber and multi-threshold models.

## DISCUSSION

In this study, we investigated and developed different models of the auditory periphery, and assessed their capacity in providing the input to an encoding model of the responses of auditory cortical neurons to a range of natural sounds. Not surprisingly, we found that the MSS model explains ferret cortical data well, since this model uses a cochlear filterbank and involves various filtering steps and non-linearities that are based on experimental observations of the auditory periphery. What is surprising, however, is that the simple cochlear models based on spectrograms, particularly the spec-Hill model, perform similarly to the MSS model. This suggests that the Spec-Hill model might capture to a large degree the functional transformation resulting from the complex processing stages of the MSS model. This holds for both single fiber and multi-fiber versions of the models.

At this point, it is reasonable to question to what extent the MSS model accurately represents the processing that takes place in the ferret auditory periphery. Although relatively few such studies have been carried out in this species, estimates of cochlear frequency selectivity in ferrets [47,48] are comparable to those made for other mammalian species, notably guinea pigs and cats, that are also commonly used in auditory research. The MSS model is derived from guinea pig data [6,7,11,12]. Recent work suggests that ferret cochlear tuning is more similar to that of guinea pigs than cats [47].

The Spec-Hill model performed similarly well, suggesting that cortical responses to natural sounds can be explained without including all the detailed properties that originate in the cochlea. One important example is phase information arising from the bidirectional displacement of hair cell stereocilia and the resulting variation in the probability of mechanoelectrical transduction channel opening with the phase of the stimulus. Spectrogram-based models throw away the fine-structure phase information of the sound waveform, while filterbank-based models do not do this to the same degree. The retention of fine-structure phase information might not be critical for the auditory cortex, as most cortical neurons do not phase lock to the waveform of sound. For example, in guinea pig primary auditory cortex, only a very small proportion of neurons (∼3%) phase lock to the fine-structure of low-frequency pure tones (<250 kHz), and most auditory cortical neurons are relatively invariant to fine-structure phase in their responses, particularly at higher frequencies [49–51]. Although the filterbank-based models may provide more biologically realistic representations of cochlear processing, retention of phase information might not be important for the stimulus selectivity of cortical neurons.

Our results also show that the choice of cochlear model can greatly influence the prediction performance of an encoding model of cortical neurons. Indeed, the observed changes in prediction performance with cochlear model were similar in size to those resulting from inclusion of other features such as gain control [28] and network structure [17,19] in models of cortical neurons. The choice of cochlear model is therefore likely to be an important factor in accounting for differences in prediction performance of LN models reported by different groups [18,20,41]. Our findings should provide a useful guide for future studies of encoding in the auditory system, suggesting which cochlear models are most suitable to use as inputs for LN models and other more complex neural encoding models.

Although the best model for each neuron operates with 64 or 128 frequency channels in the cochleagram (Supplementary Fig. 4.2), we found that increasing the number of frequency channels only moderately improves the average prediction performance of a model for the whole population of recorded neurons (Fig 4). In fact, most of the models performed relatively well with very few (<16) frequency channels. This might be because the natural sound clips that we used here have limited spectral variation over different frequency channels.

One caveat to bear in mind is that the choice of stimulus set may have some influence on the results. While we aimed to have a diverse and representative stimulus set spanning the range of natural sounds, some of which included multiple sound sources or small amounts of reverberation, this does not, of course, represent the full space of natural sounds. In particular, spatial hearing cues were not included, and reverberation and sound mixtures only present only to a very limited extent. More diverse datasets may help distinguish the models further in their capacity to predict cortical responses to behaviorally-relevant sounds.

In summary, although extensive processing takes place in the cochlea and brainstem [52], the results we present suggest that, at least under some conditions, the cortex receives a simpler functional transformation of the inputs than these properties suggest. Implementing comparable algorithms based on the key operations performed by the auditory periphery may also be useful for developing improved encoding models of central responses in other sensory modalities, such as vision and olfaction.

## MATERIALS AND METHODS

### The dataset: stimuli and neural response

The experimental data used in this study have previously been published [19] and are publicly available at https://osf.io/ayw2p/. All data were obtained from experiments performed under license from the UK Home Office and approved by the University of Oxford Committee on Animal Care and Ethical Review. They were obtained using 16- or 32-channel silicon probe electrodes (Neuronexus Technologies) from anesthetized ferrets (ketamine, 5 mg/kg/h; medetomidine, 0.022 mg/kg/h). *In vivo* extracellular recordings were made from the primary auditory cortical areas in response to *n* = 1-20 natural sound snippets (speech, ferret vocalizations, other animal vocalizations, and environmental sounds), each 5 s in duration. Each snippet was repeated 20 times. Details about the stimuli and neural responses can be found in reference [19]. In total, the dataset includes 549 single and multi-unit recordings from six adult pigmented ferrets (five female and one male), among which 284 are single units. Seventy-three of the single units had noise ratios [27,28] to the stimulus set of < 40, and these were the units used in this study. For each unit, the number of spikes was counted in 4-ms time bins and averaged over all trials. The trial-averaged neural response to stimulus *n* is henceforth denoted as *v*_*n*_(*t*), where *t* indicates the time bin. Because the beginning of a stimulus presentation often promotes various stimulus specific adaptive processes, the first 800 ms of the neural response was clipped from the data.

### Cochlear models

#### WSR model

This model was originally proposed by Wang and Shamma and was later described by Powen Ru (Matlab codes that we are using were adapted from https://github.com/tel/NSLtools) [3–5,25]. We refer to it as the WSR (Wang Shamma Ru) model after the names of the developers. It makes use of a series of filters, whose center frequencies are spaced logarithmically, followed by a sigmoid compression of the filter outputs. The sigmoid compressed outputs were then passed through a lowpass filtering stage, a lateral inhibition stage (between neighboring frequency channels), and a temporal integration stage. The source of these parameters is not strictly physiological, rather, they are abstracted from animal experiments to match perceptual processing. The filterbank of this model comprised a set of 129 given frequency channels. To make this model comparable to other models, appropriately spaced and sized subsets of the frequency channels were selected.

#### Lyon model

This model uses a cascade of filters with half-wave rectification and adaptive gain control whose center frequencies are spaced according to a logarithmic scale, except for low frequencies, where the spacing is linear, to simulate the behavior of the cochlea [2,10]. The parameters of the filters have been largely obtained from human psychophysics experiments. We adapted this model from: https://github.com/google/carfac. This implementation has several free parameters, allowing for flexible choice of spacing and bandwidth of each frequency channel in the filterbank – we set these parameters to values that would generate the desired number of frequency channels between 0.5 Hz and 20 kHz.

#### BEZ model

This is a phenomenological model of cat auditory nerve fibers [14,15,24], which we refer to as the BEZ (Bruce Erfani Zilani) model after the name of its developers (we adapted this model from: https://www.urmc.rochester.edu/MediaLibraries/URMCMedia/labs/carney-lab/codes/UR_EAR_v2_1.zip). This model has various processing stages to match the processing stages in the cochlea, the synapses with the auditory nerve and the spiking properties of the auditory nerve. Sound input is first processed through a middle ear filter, followed by a series of filters and non-linearity to account for inner hair cell and outer hair cell properties, the output of the hair cells is then processed through a non-linearity to account for synaptic release and uptake of neurotransmitter and spiking of the auditory nerve. Because the spiking is a stochastic process, for each center frequency, each of three auditory nerve fiber types was allowed to spike 20 times and the average was taken over these 20 trials. After that, we took an average over different fiber types weighted by the ratio of each fiber types in the nerve fiber population.

#### MSS model

Different components of this model have been incorporated and refined over the years; we call it the MSS (Meddis Sumner Steadman) model after the name of its developers (we adapted this model from: https://zenodo.org/record/1345757#.XW4XtXt7lPY). The MSS model is a biologically-detailed model of inner hair cells in the cochlea and the auditory nerve [6,7,11–13,16], the parameters of which are based on measurements from the guinea pig [13,16]. The MSS model includes several stages of processing. The first stage – an outer and middle ear (OME) model – uses a series of bandpass, highpass and lowpass filters to mimic stapes motion at the oval window. The second stage, the frequency decomposition stage, uses a dual resonance non-linear filter (DRNL) to model the properties of the basilar membrane. DRNL consists of two parallel pathways comprising a series of bandpass and lowpass filters, one with a linear gain and the other without a linear gain. The outputs of each parallel pathway is then combined to find the velocity of the basilar membrane displacement. Finally, transduction by the inner hair cells is modelled using a differential equation for their membrane potential, the synapse is modelled using probabilistic release of neurotransmitter and the activity of the auditory nerve (AN) fibers involves refractoriness, with the spiking of the AN fibers providing the output (Fig 1A-C). We set the center frequency of the non-linear filterbank to be log spaced between 508 Hz and 19,912 Hz. Finally, we resampled the output into 4-ms time-bins to find a windowed time-frequency representation.

#### Spec-log model

A spectrogram was produced from the sound waveform by taking the amplitude spectrum using 8-ms Hanning windows, overlapping by 4 ms. The amplitude of adjacent log-spaced frequency channels was summed using overlapping triangular windows (using code adapted from melbank.m, http://www.ee.ic.ac.uk/hp/staff/dmb/voicebox/voicebox.html) containing frequency channels ranging from 508 Hz to 19,912 Hz centre frequencies. The amplitude in each time-frequency bin was converted to log values and any value below a threshold (−40) was set to that threshold.

#### Spec-log1plus model

This model is the same as the spec-log model but uses *log(1+x)* as the compression function instead of *log(x)*.

#### Spec-power model

This model too is the same as the spec-log model, but takes the square of the spectrogram, resulting in a power spectrogram instead of an amplitude spectrogram.

#### Spec-Hill model

This model is an extension of the spec-power model where the output of spec-power model is further compressed using a non-linear Hill function[46],

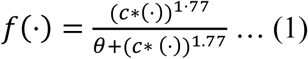

Here, c is the scaling factor, which was set to 0.01 and *θ* is the saturation parameter of the Hill function, which was set to 0.16. These values of *c* and *θ* are chosen by cross-validation on the validation set (see below).

### Multi-threshold cochlear model

The multi-threshold version of the spec-Hill model was constructed by taking the output of log compression in a spec-power model and thresholding it using values of 0 10 and 25 dB [45], and then compressing it with a Hill function with different saturation levels. Saturation was achieved by using Equation 1 with various *θ* values: 3×10^−5^, 5×10^−5^, 8×10^−5^ and c = 1×10^−4^. These values were chosen by cross-validation (see below).

### Time-lagged matrix of the normalized cochleagram

The output of each cochlear model is called a cochleagram: the frequency-decomposed transformation of sound binned into 4-ms windows. To provide input for an encoding model of an auditory cortical neuron, the cochleagrams were normalized to zero mean unit variance and for each snippet *n*, for every time *t*, a time-lagged matrix was extracted from the cochleagram. This was done according to the equation, *x*_*fqn*_(*t*) = *K*_*fn*_(*t* − *q* + 1), where *x* is the input vector, *f* is frequency, *q* is the time lag, which goes from q=1 to the maximum time lag Q, and *K*_*f*_(*t*) is the cochleagram. The resulting tensor *x*_*fqn*_(*t*) is the input to the linear nonlinear (LN) stage of the encoding scheme. Here, *Q* =50, which is 200 ms.

### The LN model

The LN model consists of a linear stage, the spectrotemporal receptive field (STRF), and a non-linear stage. The linear part of the model is:

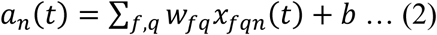

where w is a vector of the input weights and b is the background activity of the neuron. Both w and b are free parameters of the model, which were estimated by linear regression of the neuron’s firing rate *v*_*n*_(*t*) on the cochleagram tensor *x*_*fqn*_(*t*) using glmnet [43], where *a*_*n*_(*t*) is the linear estimate of *v*_*n*_(*t*). To overcome overfitting, the parameters were regularized using L1-norm (LASSO) regularization of the weights. A regularization hyperparameter λ was used to control the strength of the regularization, which was chosen by a crossvalidation procedure (see below).

The second stage of the model was a logistic activation function (sigmoid nonlinearity), which was fitted after fitting the STRF. This function is given by:

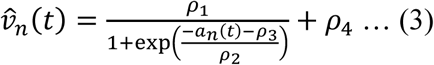

The four parameters *ρ*_*i*_ of the function were fitted by minimizing the squared error between the nonlinear estimate of the firing rate 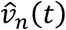 and measured firing rate *v*_*n*_(*t*).

### Cross-validation and testing of the LN model

From the data for the 20 sound stimuli, 4 were chosen as a test set which was not used during training and cross-validation. The cross-validation set (the remaining 16 stimuli) was used to fit the models using *k*-fold cross validation, where *k* = 8. The cross-validation set was randomly divided into a training set of 14 stimuli and a validation set of 2 stimuli. The LN model was trained on the training set for 18 different values of the hyperparameter λ. A log spaced range of lambda values was used, but with a somewhat lower density at the extremes. The exact values of λ used were: 1.00 × 10^−1^, 2.00 × 10^−2^, 1.17 × 10^−2^, 6.84 × 10^−3^, 4.00 × 10^−3^, 2.34 × 10^−3^, 1.37 × 10^−3^, 8.00 × 10^−4^, 4.68 × 10^−4^, 2.74 × 10^−4^, 1.60 × 10^−4^, 9.36 × 10^−5^, 5.41 × 10^−5^, 3.20 × 10^−5^, 6.40 × 10^−6^, 1.28 × 10^−6^, 2.56 × 10^−7^, and 5.12 × 10^−8^. For each LN model fitted with different λ, neural responses were then predicted for the validation set, and the correlation coefficient between the actual neural responses and the prediction was measured. This process was repeated 8 times for different non-overlapping validation sets. The model was then retrained with the whole cross-validation set using the λ value that provided the highest mean correlation coefficient over all 8 folds. Next, the retrained model was used to predict the neural responses to the test set. All the correlation coefficients and normalized correlation coefficients shown are for this held-out test set, and all the LN model parameters shown are for the retrained model.

## ACKNOWLEDGEMENTS

B. D. B. W., N. S. H., and A. J. K were supported by Wellcome Trust funding (WT108369/Z/2015/Z). M.R. was supported by the Clarendon Fund Scholarship.

## SUPPLEMENTARY FIGURES AND TABLE

**Fig S2.1:**
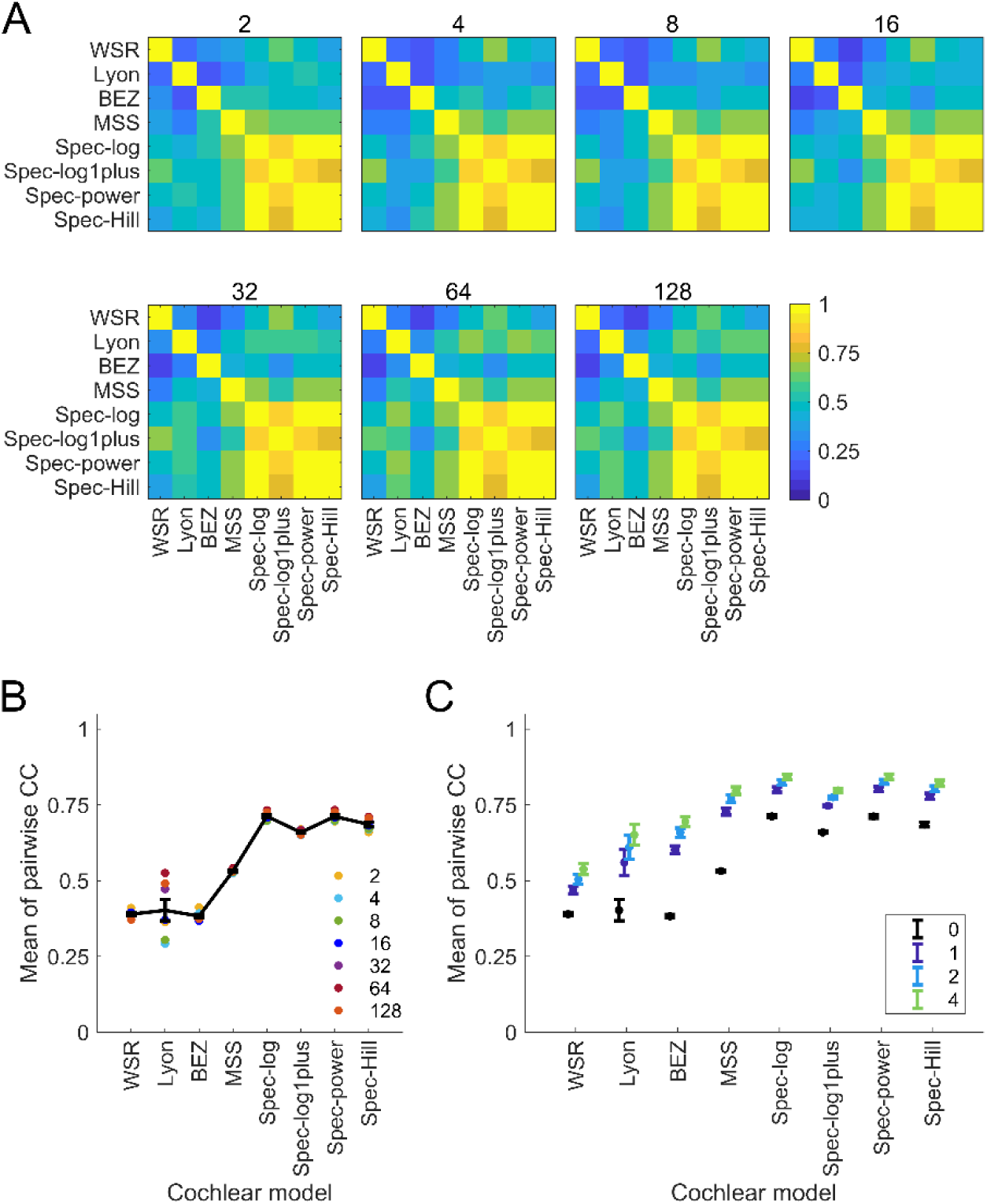
Comparison of cochleagrams produced by different models. A. Correlation-coefficient (CC) between cochleagrams produced by all possible pairs of models. The number on top of each plot shows the number of frequency channels in the cochleagram. B. Mean of the pair-wise CC of a model with the rest of the models. Colored dots show the mean for a model with a specified number of frequency channels and the black line with error bars show the mean over the means for different frequency channel numbers. C. Mean of the means of the pair-wise CCs after introducing Gaussian blurring to each cochleagram to account for spectral and temporal shift.

**Fig S3.1:**
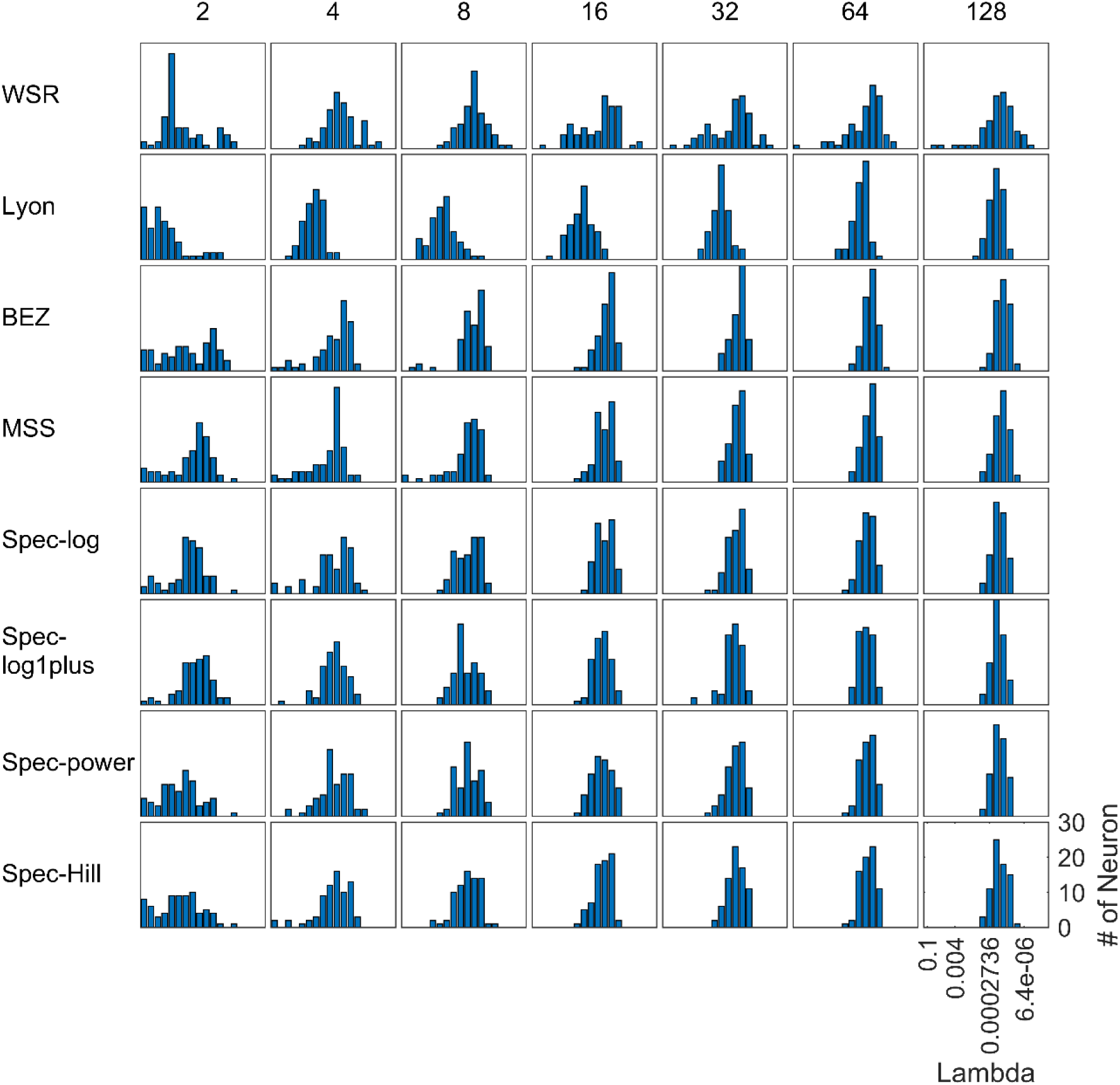
Distribution of the values of hyperparameter lambda in the LN model of each neuron. Each row shows the cochlear model that was used to generate the input for the LN model and each column shows the number of frequency channels in the cochleagram input.

**Fig S3.2:**
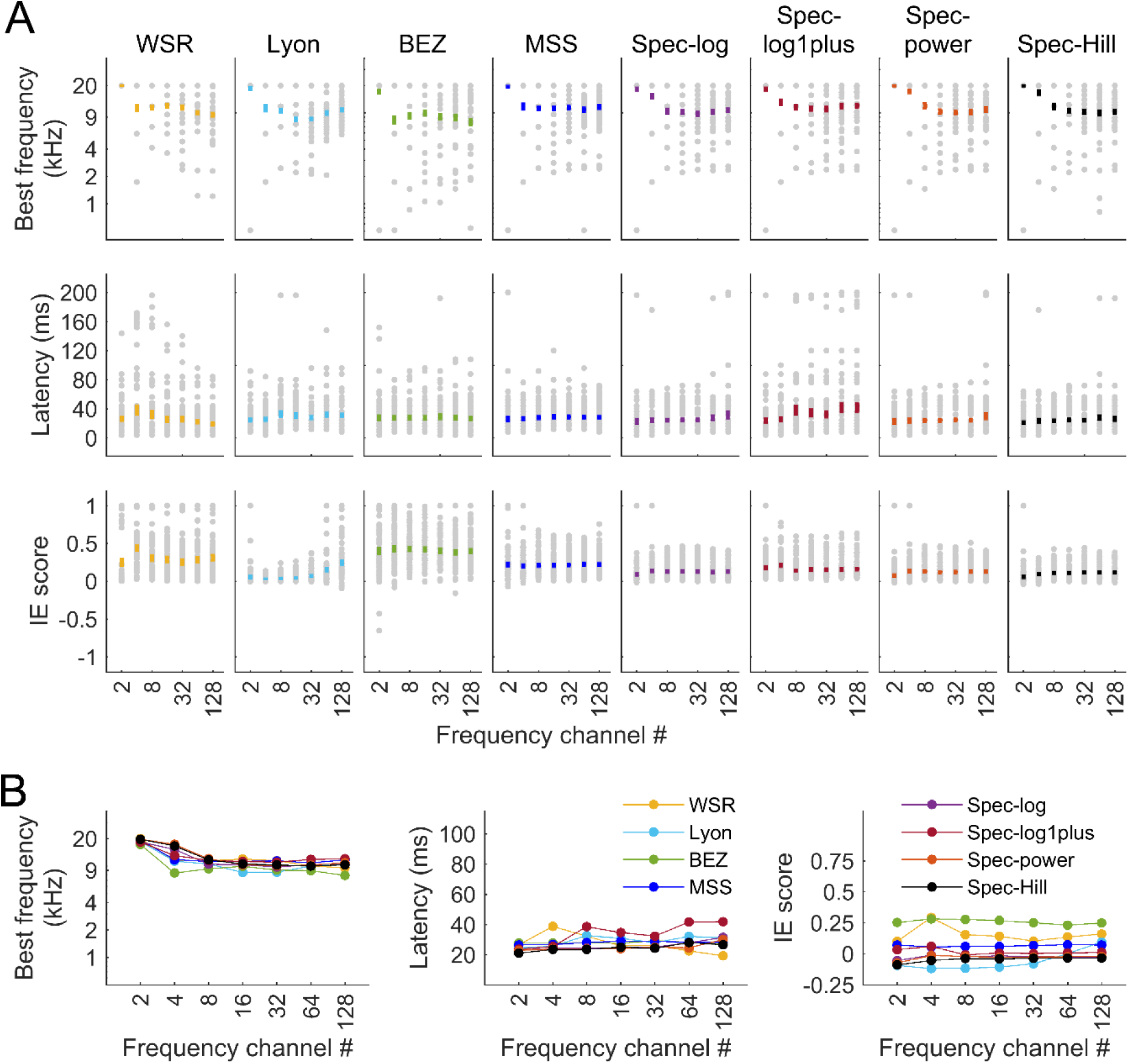
Cortical response properties estimated by using different cochleagram inputs to the LN model. A. Best frequency, latency and IE score are shown in the top, middle and bottom rows, respectively, with each plot showing the estimated values using the model specified at the top and the frequency channel specified on the x-axis. Each gray dot is one neuron and the color bar indicates the mean and the squared error of the mean. B. Mean best frequency, latency and IE score over all neurons for each model and frequency channel number.

**Fig S3.3.**
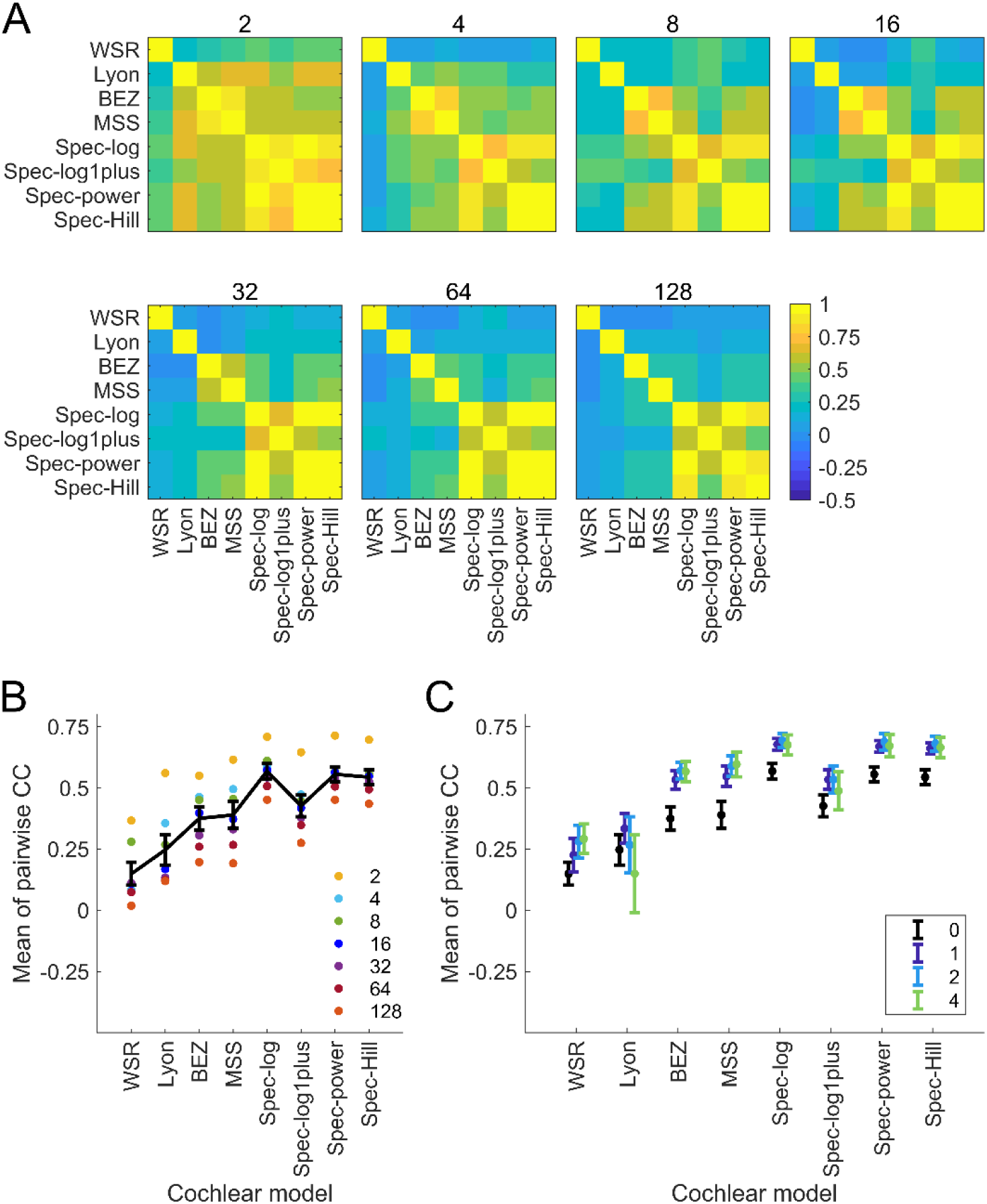
Comparison of STRFs estimated using with different cochleagram inputs. A. Correlation coefficient (CC) between STRFs produced by all possible pairs of cochlear models. The number on top of each plot shows the number of frequency channels in the cochleagram. B. Mean of the pair-wise CC of a model with the rest of the models. Colored dots show the mean for a model with a specified number of frequency channels and the black line with error bars shows the mean over the means for different numbers of frequency channels. C. Mean of the means of the pair-wise CCs after introducing Gaussian blurring to each STRF to account for spectral and temporal shift.

**Table S4.1:** Mean ± SE CC_norm_ between cortical responses and predicted responses for each model with different numbers of frequency channels. Numbers in bold indicate the top performing models.

| Models | Number of frequency channels |  |  |  |  |  |  |
| --- | --- | --- | --- | --- | --- | --- | --- |
|  | 2 | 4 | 8 | 16 | 32 | 64 | 128 |
| **WSR** | 0.301 ± 0.013 | 0.390 ± 0.013 | 0.462 ± 0.014 | 0.423 ± 0.015 | 0.429 ± 0.017 | 0.426 ± 0.016 | 0.430 ± 0.016 |
| **Lyon** | 0.320 ± 0.013 | 0.419 ± 0.012 | 0.485 ± 0.013 | 0.552 ± 0.012 | 0.614 ± 0.012 | 0.662 ± 0.013 | 0.660 ± 0.014 |
| **BEZ** | 0.313 ± 0.013 | 0.473 ± 0.011 | 0.613 ± 0.013 | 0.627 ± 0.012 | 0.635 ± 0.012 | 0.643 ± 0.012 | 0.644 ± 0.013 |
| **MSS** | 0.451 ± 0.015 | 0.580 ± 0.012 | 0.687 ± 0.012 | 0.708 ± 0.012 | 0.719 ± 0.012 | 0.723 ± 0.012 | **0.725 ± 0.012** |
| **Spec-log** | 0.670 ± 0.014 | 0.685 ± 0.014 | 0.701 ± 0.014 | 0.719 ± 0.014 | 0.715 ± 0.014 | 0.721 ± 0.014 | 0.718 ± 0.014 |
| **Spec-log1plus** | 0.562 ± 0.013 | 0.584 ± 0.013 | 0.617 ± 0.012 | 0.630 ± 0.013 | 0.627 ± 0.014 | 0.630 ± 0.013 | 0.629 ± 0.013 |
| **Spec-power** | 0.556 ± 0.013 | 0.690 ± 0.014 | 0.690 ± 0.015 | 0.715 ± 0.014 | 0.718 ± 0.014 | 0.722 ± 0.014 | 0.720 ± 0.014 |
| **Spec-Hill** | 0.542 ± 0.014 | 0.694 ± 0.015 | 0.701 ± 0.015 | 0.719 ± 0.014 | 0.720 ± 0.014 | **0.726 ± 0.014** | 0.722 ± 0.014 |

**Fig S4.2:**
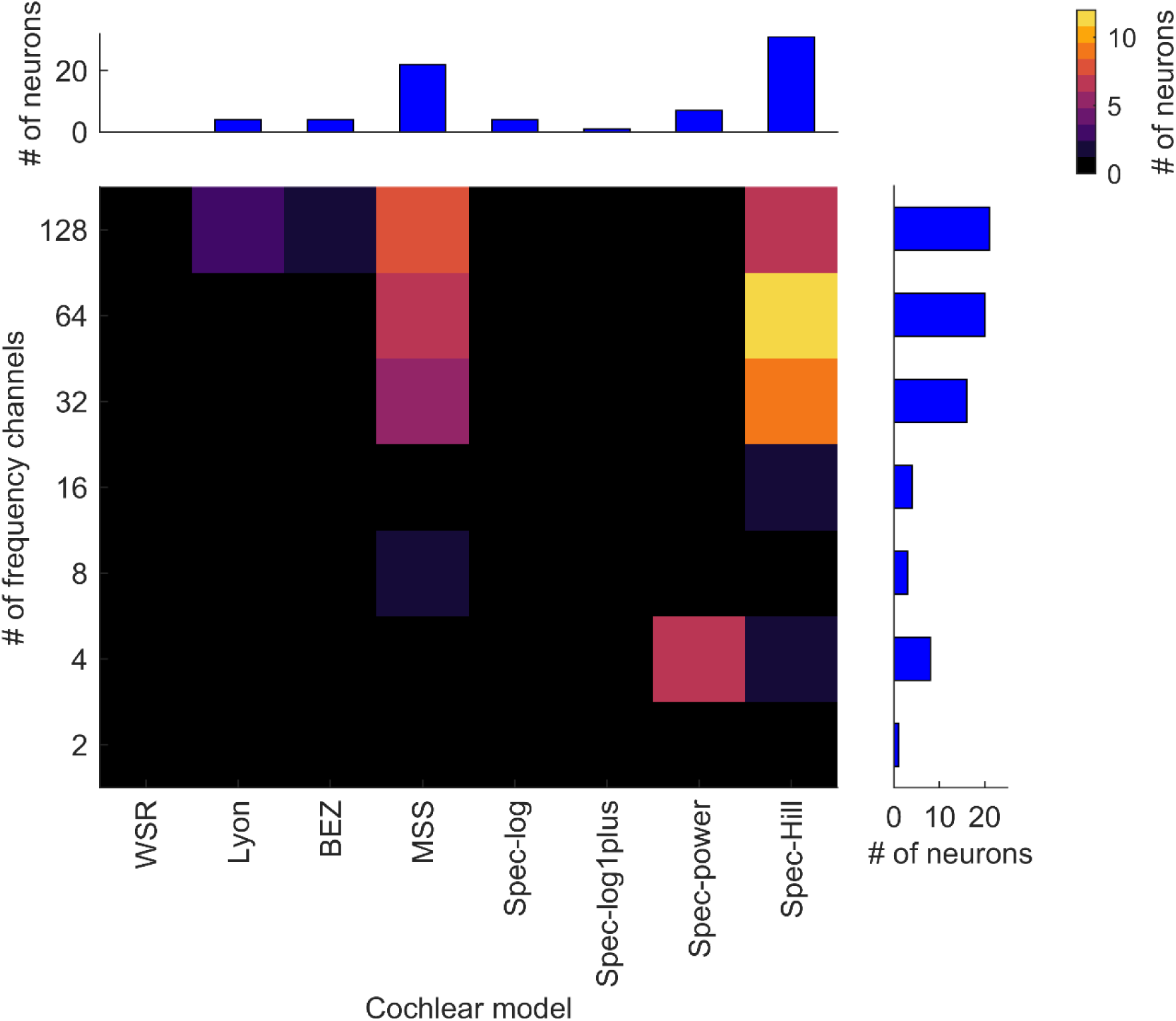
Best performing cochlear models for 73 primary auditory cortical neurons. The heatmap shows the number of neurons for which each model with a specific number of frequency channels is the best performing model and the bars show the data collapsed along each axis.

**Fig S4.3:**
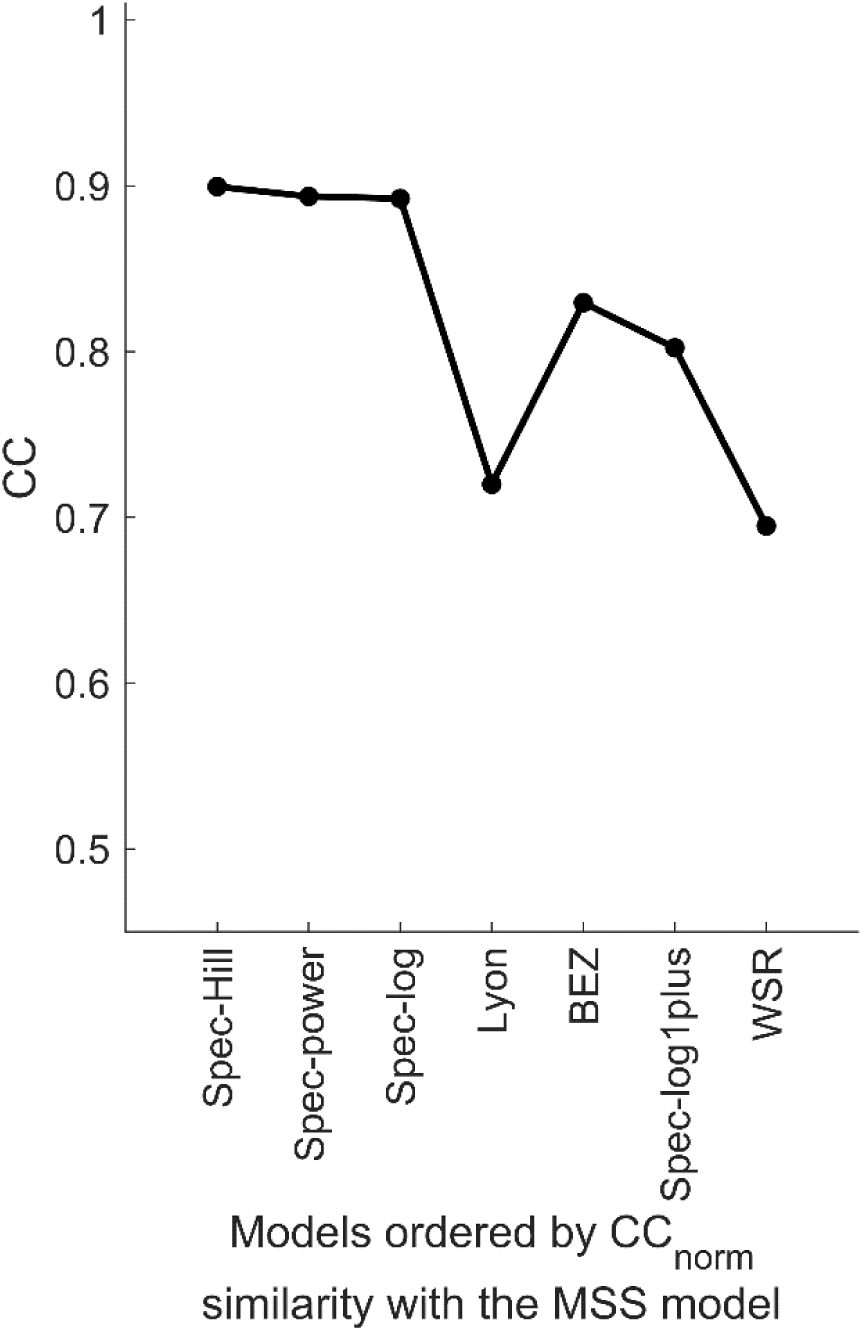
Similarity of the predicted response of the MSS model to the predicted response of other models. Each dot represents the correlation co-efficient between the response of the MSS model and the specified model in the x-axis. Models in x-axis are ordered in similarity of their CC_norm_ performance to show the relationship between prediction performance and response similarity.

## Notes

**Conflict of interest**: The authors declare no competing financial interests.

